# Intermittent theta burst stimulation at personalized targets reduces the functional connectivity of the default mode network in healthy subjects

**DOI:** 10.1101/646265

**Authors:** Aditya Singh, Tracy Erwin-Grabner, Grant Sutcliffe, Walter Paulus, Peter Dechent, Andrea Antal, Roberto Goya-Maldonado

**Affiliations:** Systems Neuroscience and Imaging in Psychiatry, Department of Psychiatry and Psychotherapy of the University Medical Center Göttingen; Department of Clinical Neurophysiology of the University Medical Center Göttingen; Core facility ‘MR-Research in Neurology and Psychiatry’, Department of Cognitive Neurology of the University Medical Center Göttingen

**Author notes:** To whom correspondence should be addressed: Dr. Roberto Goya-Maldonado.

## Abstract

Understanding the mechanisms by which transcranial magnetic stimulation protocols exert changes in the default mode network (DMN) is paramount to develop therapeutically more effective approaches in the future. A full session (3000 pulses) of 10 Hz repetitive transcranial magnetic stimulation (HF-rTMS) reduces the functional connectivity (FC) of the DMN and the subgenual anterior cingulate cortex but current understanding of the effects of a single session of intermittent theta burst stimulation (iTBS) on the DMN in healthy subjects is limited. To reduce the effects of inter-individual variability in functional architectures, we used a novel personalized target selection approach based on each subject’s resting state fMRI for an unprecedented investigation into the effects of a single session (1800 pulses) of iTBS over the DMN in healthy controls. 26 healthy subjects participated in a double-blind, crossover, sham-controlled study. After iTBS to the personalized left dorsolateral prefrontal cortex (DLPFC) targets, we investigated the time lapse of effects in the DMN and its relationship to the harm avoidance (HA) personality trait measure (Temperament and Character Inventory/TCI). Approx. 25-30 minutes after stimulation, we observed reduced FC between the DMN and the rostral anterior cingulate cortex (rACC). About 45 minutes after stimulation the FC of rACC strongly decreased further, as did the FC of right anterior insula (rAI) with the DMN. We also report a positive correlation between the FC decrease in the rACC and the HA domain of TCI. Our results show how iTBS at personalized left-DLPFC targets reduces the FC between DMN and the rACC and rAI, regions typically described as nodes of the salience network. We find that HA scores can potentially predict iTBS response, as has been observed for HF-rTMS.

## Introduction

Both the large variability of responses in the treatment of depression by the FDA-approved 10 Hz repetitive transcranial magnetic stimulation (rTMS) protocol has led to a world-wide demand for better techniques or improved protocols. The non-inferior antidepressant efficacy of the 3 min/session theta burst protocol^1^ compared to 37.5 min/sessions of conventional 10 Hz rTMS protocol has played a role in increasing the use of the theta burst protocol for antidepressant treatment^2–5^. The connectivity changes underlying the effects of intermittent theta burst stimulation (iTBS) delivered at the left dorsolateral prefrontal cortex (DLPFC) remain unexplored. Many factors contribute to inter-individual variability, including natural variation in anatomy and functional connectivity. Here we use a previously validated target selection method to improve precision of coil localization and investigated the effects of iTBS on the relevant brain networks that cover the left DLPFC and the anterior cingulate cortex (ACC).

The TBS protocol was developed to mimic rodent^6,7^ and human hippocampal activity^8^, where a combination of gamma-frequency spike patterns superimposed on theta rhythms^9^ was found. It involves application of a burst of three TMS pulses every 20 milliseconds (50 Hz), which is repeated five times per second (5 Hz)^10,11^. When delivered continuously (continuous TBS – cTBS) for 40 seconds, it results in reduced corticospinal excitability, while when administered in an intermittent fashion (iTBS) it results in increased corticospinal excitability ^9^. Studies of TBS stimulation on motor cortex have shown plasticity changes beyond the duration of stimulation typically lasting in the range of 30 minutes^11,12^.

Beyond local effects under the stimulation coil, plasticity changes in brain’s altered functional connectivity away from stimulation point e.g. the DLPFC^13^ are likely relevant to the treatment of psychiatric disorders, which has been suggested to result from aberrant brain functional connectivity^14^. The DMN, consisting of the medial prefrontal cortex, posterior cingulate cortex and areas of posterior parietal cortex ^15^, is usually hyperconnected to subgenual ACC (sgACC) in depression ^15,16^. A reduction of this hyperconnectivity has been related to a reduction of symptoms ^17^. Using 10 Hz rTMS as antidepressant treatment, a study has recently replicated the prediction of symptomatic alleviation in depression when aberrant sgACC connectivity with the DMN is decreased, which happened in responders but not in non-responders^18^. Furthermore, such effects over networks in healthy subjects have been shown in our previous work ^19^. Already after a single session of 10 Hz rTMS (3000 pulses), delivered at personalized left DLPFC sites, a reduction in the connectivity between the sgACC and the DMN was evidenced, most strongly in subjects with lower harm avoidance (HA) scores from the Temperament and Character Inventory (TCI).

Given the central involvement of the DMN in the pathophysiology of depression and the importance of a shorter protocol such as iTBS in reducing symptoms,^15,20–31^ here we aimed to uncover if a single session of a prolonged iTBS protocol (1800 pulses) in healthy subjects would result in reduced DMN connectivity to the ACC. We applied a single session of iTBS at personalized left DLPFC sites and analyzed the DMN during three time windows after stimulation in a double-blind, crossover, and sham-controlled study. Given that the nature of iTBS differs from 10 Hz rTMS, we were interested in whether the modulation across the sgACC is similar. Moreover, based on a negative correlation seen between HA scores and coupling changes of the sgACC and the DMN after one session of 10 Hz rTMS^19^, we hypothesized a correlation between iTBS induced changes in DMN and HA scores.

## Materials and Methods

### Participants

Healthy subjects ages 18-65 were enrolled in the study. We evaluated the subjects with structured clinical interviews and ruled out current or prior psychiatric disorders. We performed the experiments in agreement with relevant guidelines and regulations^32,33^. The Ethics Committee of the University of Medical Center Göttingen approved the study protocol and subjects provided their informed consent before investigation.

### Study design

The study reported here with healthy subjects is a sham-controlled, double-blind (subject and interviewer), crossover study. We conducted the experiments over three sessions (each session on a different day, Figure 1) with each session separated by at least one week.

**Figure 1:**
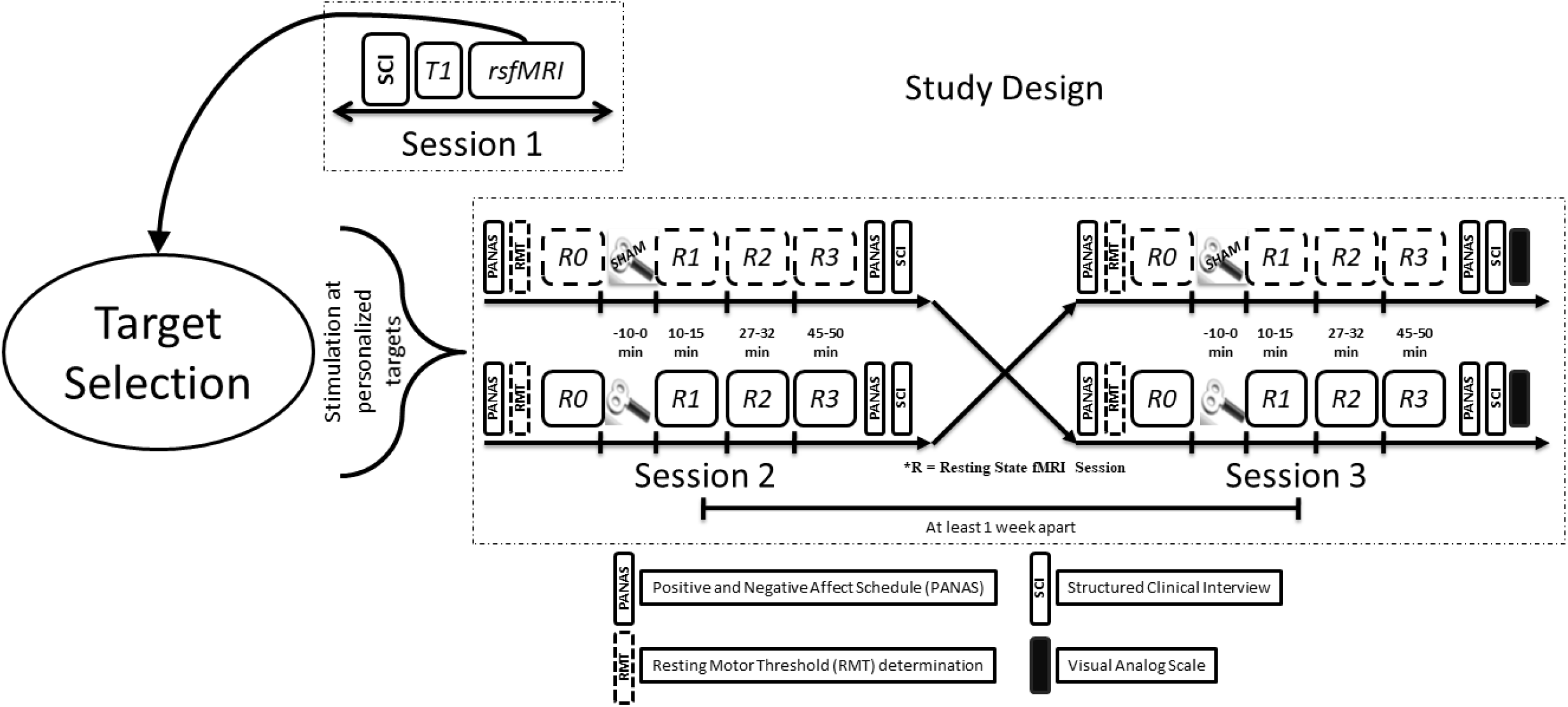
A schematic representation of the study design. In session 1, we obtained the informed consent and collected the information from Structured Clinical Interviews (SCI). After this we acquired a structural (T1-weighted) and functional (rsfMRI) image. During rsfMRI, the subjects were instructed to fixate at a ‘+’ and mind wander, while their open eyes were monitored by eye tracking. Personalized target were found using each subject’s rsfMRI as has been described elsewhere^19^. Using online neuronavigation, we delivered real or sham iTBS, in a counterbalanced and pseudo-randomized fashion, at 80% of resting motor threshold. Baseline and three post-iTBS rsfMRI scans were acquired. The subjects also completed the Positive and Negative Affect Schedule (PANAS) both before and after the sessions, and a visual analog scale (VAS) for perceived effects of iTBS on mental state and scalp sensation at the end of the sessions.

### Session 1

In session 1, the interviewer administered a Structured Clinical Interview (SCI) consisting of the Beck Depression Inventory II (BDI II), Montgomery-Asberg Depression Rating Scale (MADRS), Hamilton Depression Rating Scale (HAM-D) and Young Mania Rating Scale (YMRS). In addition to the SCI, to further establish the mental health and well-being of the subjects we asked them to complete the Symptoms Checklist 90-revised (SCL 90-R), Temperament and Character Inventory (TCI), Positive and Negative Syndrome Scale (PANSS), Life Orientation Test – Revised (LOT-R), Barratt Impulsiveness Scale (BIS), a handedness questionnaire^34^ and a vocabulary-based intelligence test (MWT). The interviewer obtained written informed consent from the subject after completing an evaluation of the inclusion and exclusion criteria. Next, we acquired structural T1-weighted MRI and resting state functional MRI (rsfMRI) scans for our novel method of personalized target selection (see Figure 1 for further details). The process of personalized left DLPFC target selection has been described previously^19^.

### Session 2 and Session 3

To allow wash out of any potential iTBS effects session 2 and session 3 were separated by at least a week. After determining the resting motor threshold (RMT), we applied iTBS, at 80% RMT. We navigated to the personalized left DLPFC target using an online neuronavigation system (Visor 1 software, ANT Neuro, Enschede, Netherlands). We obtained a pre-iTBS (baseline) rsfMRI scan (R0) followed by three post-iTBS rsfMRI scans. Subjects completed the Positive and Negative Affect Schedule (PANAS^35^) at the before and after the experiment on session 2 and session 3. This allowed us to follow any short-term changes in the subjects’ mood because of iTBS. Figure 1 pictorially details the study design.

### rTMS protocol

We delivered iTBS using a MagVenture X100 with Mag-option and a “figure of 8” MCF-B65 cooled butterfly coil at the targets selected using each individual subject’s rsfMRI (see ^19^). We used stimulation parameters from Li C-T et al., 2014^5^ [3 pulses burst at 50 Hz delivered at 5 Hz for 2 seconds with an 8 second inter train interval, total 60 trains delivered during 9 minutes 30 seconds]. For the sham condition, we rotated the coil by 180° along the handle axis of the coil as in our previous work^19^.

### Image Acquisition

We collected the functional and the structural (T1- and T2-weighted scans with 1-mm isotropic resolution) data with a 3T MR scanner (Magnetom TIM TRIO, Siemens Healthcare, Erlangen, Germany) using a 32-channel head coil. The T2*-weighted multi-band gradient-echo echo-planar imaging sequence provided by the Center for Magnetic Resonance Research of the University of Minnesota^36,37^ had the following parameters: repetition time of 2.5 s, echo time of 33 ms, flip angle of 70°, 60 axial slices with a multi-band factor of 3, 2×2×2 mm, FOV of 210 mm, with 10% gap between slices and posterior to anterior phase encoding. The rsfMRI data were acquired with 125 volumes in approx. 5 minutes. The gradient echo field map was acquired with repetition time of 603 ms, echo times of 4.92 ms (TE 1) and 7.38 ms (TE 2), flip angle of 60°, 62 slices, FOV of 210 mm, 2×2×2 mm, with 10% gap between slices and anterior to posterior phase encoding.

### Imaging Data Analysis

We preprocessed the individual rsfMRI data using SPM12 (http://www.fil.ion.ucl.ac.uk/spm/software/spm12/) and MATLAB (The MathWorks, Inc., Natick, MA, USA) to execute the following state-of-the-art steps: slice time correction, motion correction, gradient echo field map unwarping, normalization, and regression of motion nuisance parameters, cerebrospinal fluid and white matter. Following this, we temporally concatenated the data for group independent component analysis (ICA) with FSL 5.0.7 software^38^. We visually identified the independent component (IC) that best resembled the DMN and another IC that covered the left DLPFC (IC-DLPFC). We back reconstructed this IC representing the DMN in the normalized rsfMRI data of individual subjects, r-to-z transformed and compared across the groups using a factorial design ANOVA (Real [R0, R1, R2, R3] versus Sham [R0, R1, R2, R3]).

#### Extraction of parameter estimates (functional connectivity strengths)

We used MarsBar^39^ to extract the parameter estimates (beta weights) of the rostral anterior cingulate cortex (rACC; 5 mm radius sphere) and subgenual anterior cingulate cortex (sgACC; 5 mm radius sphere) centred on independent coordinates from a meta-analysis of functional large-scale networks in depression^40^ and our previous work on 10 Hz rTMS effects on DMN^19^, respectively. The parameter estimates for the personalized left DLPFC sites were extracted using 2 mm radius sphere ROI centered around the personalized targets, in line with our previous work^19^. Figure 2A highlights an example subject showing the IC-DLPFC (in warm color), the network from which the parameter estimates using a personalized left DLPFC target ROI (blue sphere) is extracted. Figure 2B shows all the personalized ROIs which were used for parameter estimate extraction.

**Figure 2:**
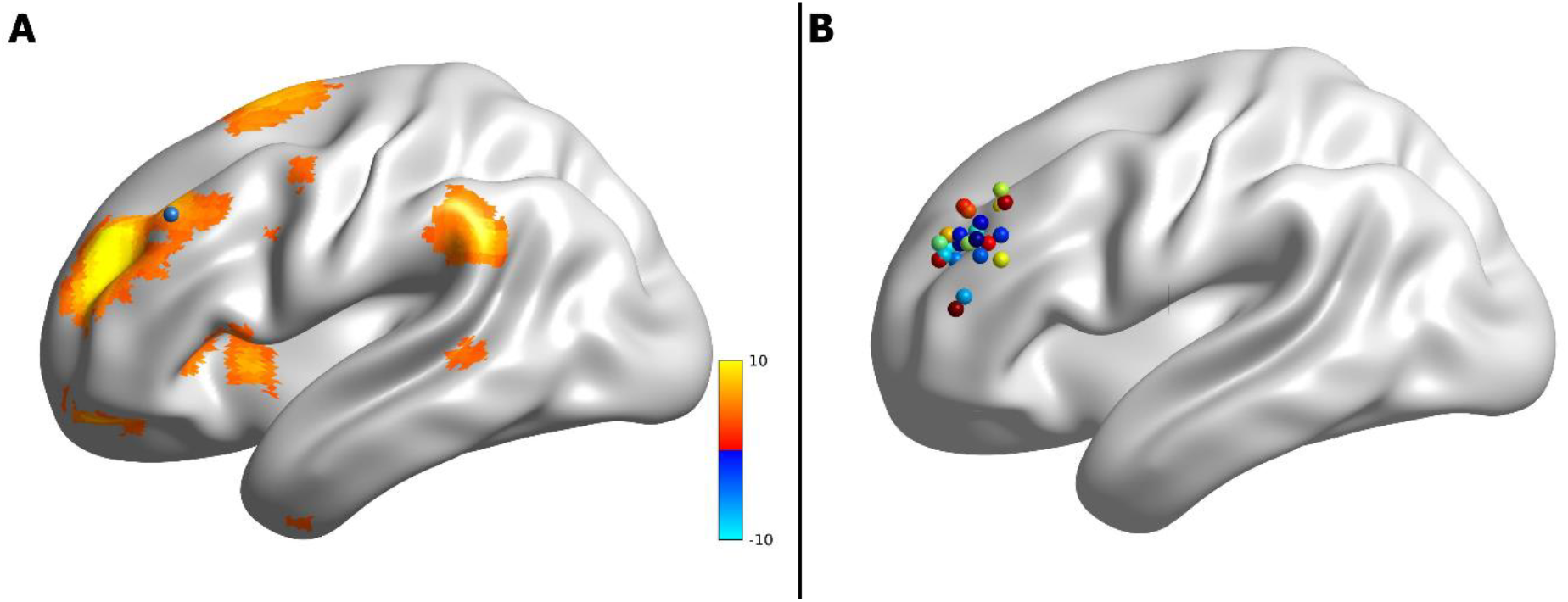
A) An example of IC-DLPFC (warm colors) for a single subject from which the parameter estimates of personalized stimulation site (blue sphere) were extracted. B) Personalized left DLPFC stimulation sites of all subjects from which parameter estimates were extracted.

### Statistical Analysis

Using a factorial design ANOVA in SPM12 we compared the time windows of rsfMRI across real and sham conditions using a, and report results surviving a statistical threshold of p<0.05 FWE whole brain corrected for multiple testing. We ran Pearson’s correlation tests between rACC functional connectivity strengths and the harm avoidance domain of the TCI using MATLAB. We used R to run two-way t-tests to compare the scores from YMRS, HAM-D, MADRS, PANAS, VAS and BDI II for real and sham stimulation sessions.

### Data availability statement

Owing to restrictions in the data sharing consent obtained from the participants of the study, the datasets generated and analysed cannot be made publicly available.

## Results

Twenty-nine healthy subjects (11 females, mean age of 28 years +/-8 years) signed up for the study. Two subjects (both females) were dropped from the study due to failure to locate their personalized left DLPFC target and one subject (male) dropped out of study due to discomfort from stimulation. Thus 26 subjects were included in final analysis, none of whom reported any adverse effects during or after stimulation.

### Functional connectivity changes after real stimulation

After a full single session of iTBS (1800 pulses) we observed reduced functional connectivity of the rACC and dorsal ACC (dACC) with the DMN, during the R2 rsfMRI session (27-32 minutes post-stimulation) when compared to R1 rsfMRI session (10-15 minutes post stimulation) (Figure 3 [A1-A2]). Even more interesting was the effect on the functional connectivity of DMN during the R3 rsfMRI (45-50 minutes post-stimulation), which increased in spatial extent. During R3, the area of significantly reduced functional connectivity of the DMN spread to include the medial prefrontal cortex (mPFC) and frontal poles, as seen in Figure 3 [B1-B2]. Additionally, the right anterior insula (AI) shows decreased functional connectivity to the DMN during R3 rsfMRI (figure 3 [B3-3B4]). These findings were not seen in the sham condition. Changes in clinical scales were neither expected nor identified. Also, it is important to note that when comparing the DMN only across real iTBS rsfMRI sessions without sham correction, we see the same regions decoupling from the DMN (Supplementary Figure 1), except by smaller mPFC and larger rAI blobs in the R2 rsfMRI. In this case, the decoupling of the right AI is more pronounced, showing significantly reduced functional connectivity even during the R2 rsfMRI.

**Figure 3:**
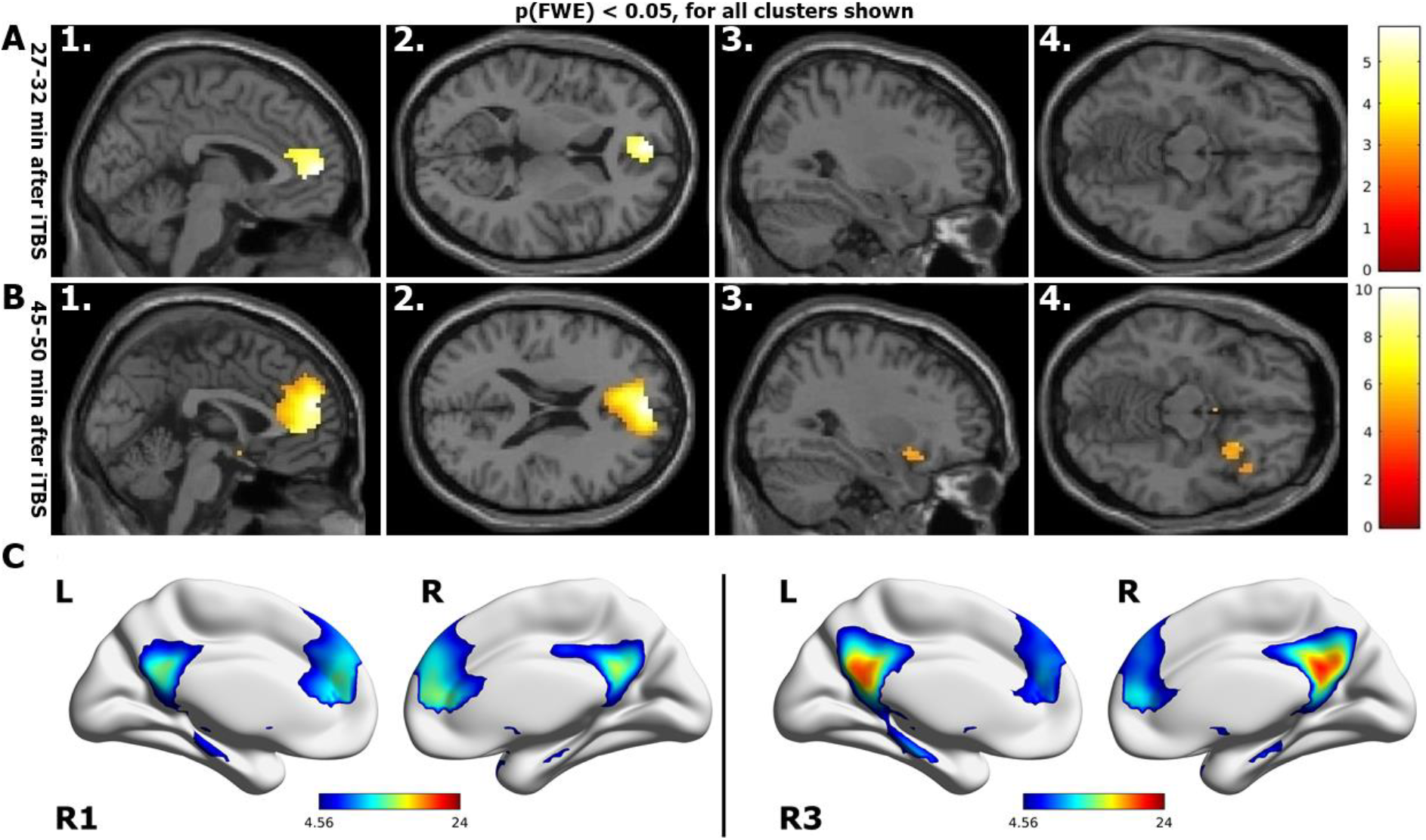
Regions that show reduced functional connectivity to DMN after stimulation (real-sham condition, whole-brain corrected p_FWE_<0.05): A1-2) About 27 minutes after iTBS, the rACC and dACC disengage from the DMN. B1-2) About 45 minutes after stimulation, the functional connectivity has further reduced, extending to the mPFC and B3-4) the right AI. C) DMN during R1 rsfMRI and R3 rsfMRI session after real stimulation.

### Functional connectivity changes in the left DLPFC and the rACC along time

To have a better understanding of the effects of iTBS, we extracted the parameter estimates of two regions of interest in the real condition: the personalized left DLPFC and the rACC. We used a spherical ROI of 2 mm radius centered at the left DLPFC target to extract its parameter estimates from the IC-DLPFC (see methods for definition of IC-DLPFC). We extracted the parameter estimates of the rACC from the DMN. Following an earlier study of the ACC with a 10 Hz protocol^19^, a spherical 5 mm radius ROI was used with coordinates obtained from an independent meta-analysis^40^. The plot (Figure 4) shows that the DMN functional connectivity of the rACC increases from the R0 to the R1 rsfMRI window. Subsequently, a functional connectivity decrease in the rACC from R1 to R2 is sustained until R3. An insignificant increase in the IC-DLPFC functional connectivity of the left DLPFC from R0 to R1 is also seen. This functional connectivity returns to a value close to baseline during R2 and increases during R3 rsfMRI. The green dotted line represents the correlation coefficients between the parameter estimates of the sgACC and the left DLPFC. It shows that as the effect of iTBS becomes more prominent, the correlation between these regions goes from negative to more and more positive, not returning to baseline within 50 minutes after iTBS. We explored the changes in functional connectivity of these regions for sham condition (Supplementary Figure 2) and observed minor changes in the median of parameter estimates (ranging between 0-0.02) and in the correlation coefficient (between 0-0.17). However, the functional connectivity fluctuates around the baseline during all rsfMRI sessions.

**Figure 4:**
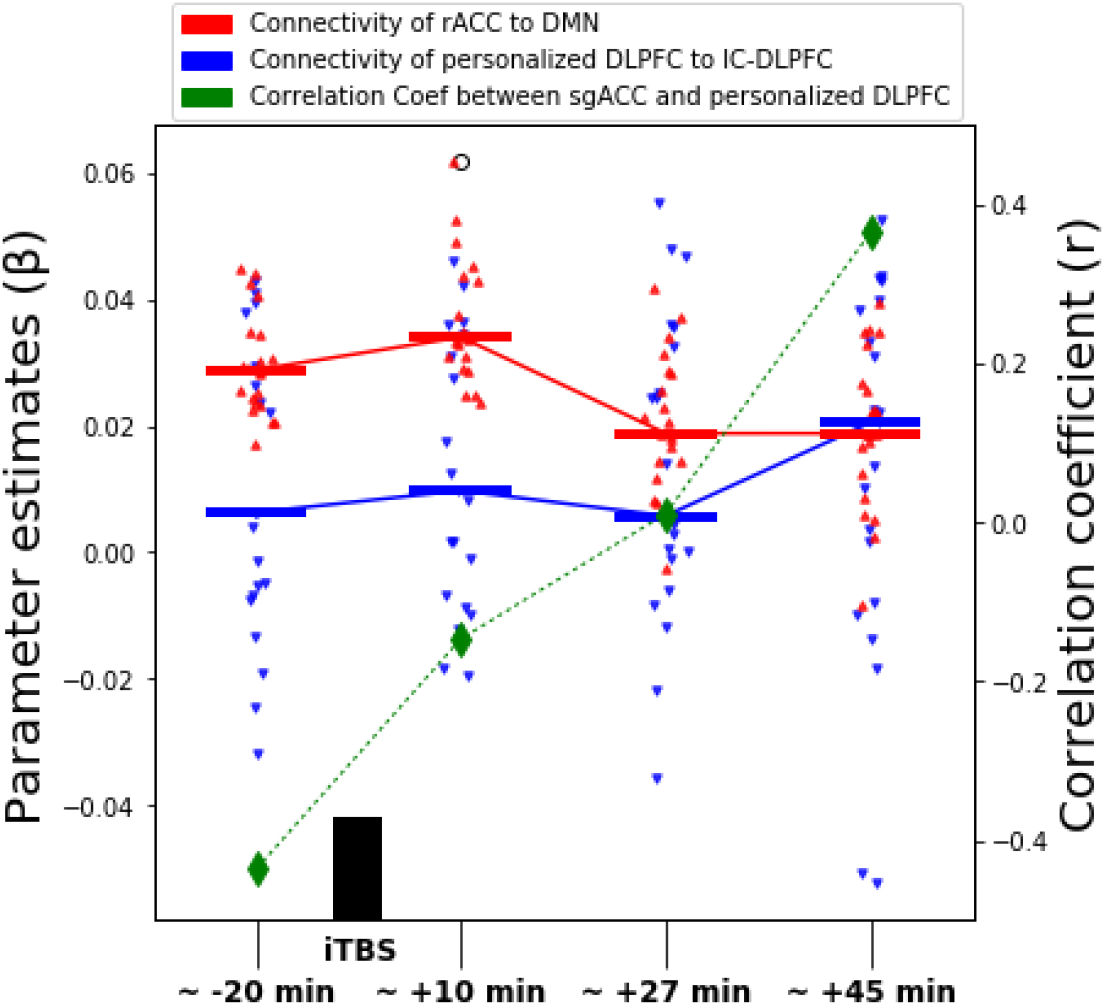
Left axis shows the parameter estimates of left DLPFC (blue) and rACC (red) of IC-DLPFC and DMN, respectively. Dots represent the individuals and horizontal lines depict the median of the parameter estimates for the respective rsfMRI window. The right axis plots the correlation coefficients between the DLPFC and the rACC, showing that the effect of a single session of iTBS progressively changes the correlation between these ROIs from negative to positive.

### Harm avoidance – a predictor of iTBS response?

Our previous work has identified a negative relationship between harm avoidance scores on TCI and the changes induced by 10 Hz rTMS in the right sgACC during R2 rsfMRI compared to R1 rsfMRI^19^. We hence explored if such a relationship existed also between the harm avoidance scores of subjects in the current study and the observed decrease in the functional connectivity of rACC during R2 rsfMRI compared to R1 rsfMRI. We identify a positive correlation between the harm avoidance measure and the decrease in functional connectivity of the rACC, only after real stimulation (r = 0.6052, p value = 0.013) but not after sham stimulation (r = −0.1233, p value = 0.6491). This indicates that the higher the harm avoidance score of the subjects the more they showed a decrease in their rACC functional connectivity to DMN (Figure 5).

**Figure 5:**
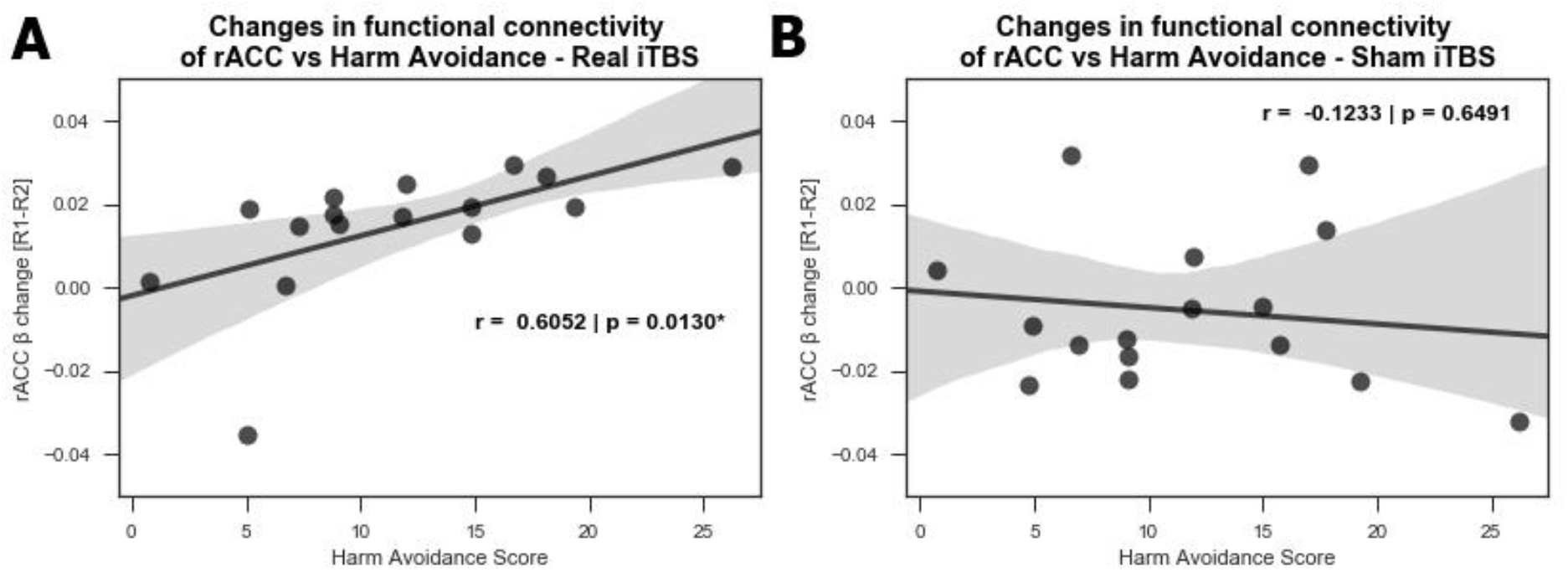
Correlation between the harm avoidance score and the changes observed in rACC during R2 rsfMRI compared to R1 rsfMR for A) real and B) sham conditions. A significant positive correlation is observed after real stimulation only.

## Discussion

In this double-blind, sham-controlled study, we have determined for the first time the connectivity changes of the DMN in the healthy brain for up to 50 minutes after iTBS (1800 pulses protocol). As expected, after personalized left DLPFC stimulation (Figure 2B) we see a decrease in the functional connectivity of the DMN, mainly with the rACC and dACC during the R2 rsfMRI window (Figure 3 A1-2, about 27-32 minutes after stimulation). This decrease is sustained in the rACC and additionally extends to the mPFC and right AI during the R3 rsfMRI window (Figure 3 B1-4, about 45-50 minutes after stimulation). In agreement with the literature^18,41^, we see at baseline a negative correlation between the parameter estimates of sgACC and the personalized left DLPFC (Figure 4, green diamond at ~20 minutes before iTBS). As the functional connectivity changes in both left DLPFC and rACC within their own networks (Figure 4, red and blue curves), the negative correlation between sgACC and left DLPFC becomes progressively positive (Figure 4, green dotted curve). Finally, we observe a positive correlation between the HA score and the connectivity changes observed with the DMN in the rACC (Figure 5A), which implies that this measure can possibly predict the magnitude of functional connectivity changes induced by iTBS in rACC.

A dynamic system known as the triple network model has been suggested to explain the fast adaptive qualities of the brain^42,43^. According to the triple network model, a task positive network corresponding to the central executive network (CEN) is active when the brain is engaged in cognitive tasks or allocating attention to external stimuli^25,44^. The DMN (aka task negative network) is active antagonistically to the task positive network, being active when resources are internally allocated during introspective thoughts or autobiographical memories^45^. A dynamic interplay between the task positive and task negative network is required to quickly reallocate resources towards internal or external stimuli according to immediate demands. It has been shown that this “circuit breaker” role is played by the salience network (SN)^43^, with the dACC and right AI as the main network nodes. We identified these regions as being decoupled from the DMN in healthy subjects after a single session of a prolonged iTBS protocol (1800 pulses) (Figure 3). Although the changes evidenced here may not directly translate to the context of psychopathology, it was very promising to see these nodes as part of our results. The AI is considered to be the essential hub of the SN because it mediates the information flow across the brain to different networks and switches between central-executive and default-mode networks^42,43,46^.

In depression, the SN shows aberrant functional connectivity to the DMN and CEN^23,25,47^. One of the network-based hypotheses of depression conjectures that the increased interaction between the SN and the DMN results in pathologically increased allocation of resources to negative information about the self, e.g. ruminative thoughts^30,48^. Considering the proven efficacy of iTBS for treatment of depression and speculating that the effects seen in healthy participants would extend to patients, the mechanism by which iTBS may initially influence the symptomatology of depression could be by “normalizing” the pathologically increased interaction between the SN and the DMN. In line with this reasoning, Iwabuchi et al.^49^ has shown in patients with depression that fronto-insular and SN connectivity interactions correlated positively with HAM-D score change at the end of a 4-week iTBS protocol. They have also described that better clinical outcomes are associated with reduced connectivity between dorsomedial prefrontal cortex (DMPFC) and bilateral insula^47^. Their results in conjunction with ours highlight the importance of investigating the AI and dACC as SN nodes involved in responsiveness to iTBS.

Using a different approach, Baeken et al. ^50^ have shown that the functional connectivity of the sgACC and medial orbitofrontal cortex (mOFC) is increased during accelerated iTBS in depression patients. Their results stem from seed-based analysis of the sgACC after iTBS. One possible reason for the discrepancy between their and our results may be the method of analysis, as seed-based analysis focuses on the functional connectivity of a predefined ROI while ICA allows exploration of functional connectivity changes of the whole brain without having to pre-define a ROI. In contrast to our previous work that identified the sgACC as the main region decoupled from the DMN after a single session of personalized 10 Hz rTMS^19^, the strongest changes in connectivity after iTBS are not with the sgACC, but rather the rACC and dACC as mentioned above. However, due to the relevance of the sgACC, we further explored the beta weights from this region and evaluated its relation to the left DLPFC at baseline and up to 50-min after stimulation. We evidenced a shift of correlation between these regions from negative to positive within the observation time (Figure 4, green dotted curve). This suggests the participation of the sgACC in the effects driven by iTBS, even though it is not directly engaged by it. The striking similarity between the red curves seen in the rACC after iTBS (Figure 4) and in the sgACC after 10 Hz TMS (Figure 5 in Singh et al.^19^) suggests that sgACC is rather the first target after 10 Hz rTMS (under standard dose of 3000 pulses). Another important aspect that might have contributed to differences between our and the results of Baeken et al.^50^ is that we stimulated functionally relevant sites within the left DLPFC, as opposed to their structural selection. Of course, the most profound difference is that our study closely evaluated connectivity changes after one session of iTBS in healthy subjects, whereas Baeken et al.^50^ evaluated patients with depression after 20 stimulation sessions. It must be considered that the complexities associated with the underlying pathophysiology of depression could have contributed to differences in how iTBS interacts with brain regions and networks. Our results shed light on other relevant regions that respond to a single session of iTBS in the healthy brain. Future work examining brain networks in patients before and after 20 iTBS treatment sessions should close these knowledge gaps.

We also evidenced a positive correlation between changes in the HA score on the TCI and the functional connectivity to the rACC in the DMN (Figure 5A). This indicates that the higher the subjects scored on the HA domain, the stronger the reduction in functional connectivity. This correlation indicates that it might be possible to utilize the HA to predict the extent of DMN-rACC coupling changes induced by iTBS. Interestingly, the correlation between connectivity changes and HA scores replicates the time window in which this was seen in an independent sample using 10 Hz rTMS^19^, although in opposite direction and in a different brain region, the sgACC. Considering the rACC and the sgACC are the regions whose connectivity to the DMN is affected by iTBS and 10 Hz TMS respectively, we speculate that HA scores may facilitate identification of participants who will present stronger DMN changes in response to TMS protocols. If this holds true for clinical samples, we hypothesize that such a measure could be used to determine beforehand who would benefit most from one stimulation protocol or the other. We hope future research in precision medicine will investigate this using different TMS protocols, considering the direct clinical application and potential relevance to improving treatment response.

There are limitations to this study. It should be noted that our choice of a single session of 1800 pulses iTBS for uncovering its effect on the DMN in healthy brains was based on ethical reasons, as applying 20 sessions of iTBS as done in patients^51^ would not be prudent. While the results presented have been controlled for using a sham stimulation, it must be noted that the method used for sham stimulation allowed some lingering current on the sham side. The strength of this current was low and not enough to elicit motor response. However, the fact remains that the sham condition used was not completely passive and hence could be interpreted as an active sham. Lastly, the diseased state of the brain, e.g. in depressive state, is likely to influence interactions between brain networks in response to multiple sessions of iTBS. Therefore, assumptions based on healthy samples must be made cautiously.

In conclusion, by means of a double-blind, sham-controlled crossover study involving healthy subjects, we show that a single session of iTBS results in decoupling of the rostral/dorsal ACC, followed by the mPFC and the right AI, with the DMN. The interaction between the personalized sites of stimulation at the left DLPFC and the rACC shows a progressive shift from negative to positive correlation. Lastly, connectivity changes in the rACC induced by a single real session of iTBS in the healthy brain positively correlated with the HA score on the TCI scale.

## Supplementary information

Supplementary information is available at Molecular Psychiatry’s website.

## Acknowledgements

This work was supported by the German Federal Ministry of Education and Research (Bundesministerium fuer Bildung und Forschung, BMBF: 01 ZX 1507, “PreNeSt - e:Med”).

## References

1 Blumberger DM, Vila-Rodriguez F, Thorpe KE, Feffer K, Noda Y, Giacobbe P et al. Effectiveness of theta burst versus high-frequency repetitive transcranial magnetic stimulation in patients with depression (THREE-D): a randomised non-inferiority trial. Lancet 2018; 391: 1683–1692.

2 Bakker N, Shahab S, Giacobbe P, Blumberger DM, Daskalakis ZJ, Kennedy SH et al. rTMS of the Dorsomedial Prefrontal Cortex for Major Depression: Safety, Tolerability, Effectiveness, and Outcome Predictors for 10 Hz Versus Intermittent Theta-burst Stimulation. Brain Stimul 2015; 8: 208–215.

3 Duprat R, Desmyter S, Rudi DR, van Heeringen K, Van den Abbeele D, Tandt H et al. Accelerated intermittent theta burst stimulation treatment in medication-resistant major depression: A fast road to remission? J Affect Disord 2016; 200: 6–14.

4 Bulteau S, Sébille V, Fayet G, Thomas-Ollivier V, Deschamps T, Bonnin-Rivalland A et al. Efficacy of intermittent Theta Burst Stimulation (iTBS) and 10-Hz high-frequency repetitive transcranial magnetic stimulation (rTMS) in treatment-resistant unipolar depression: Study protocol for a randomised controlled trial. Trials 2017; 18. doi:10.1186/s13063-016-1764-8.

5 Li C-T, Chen M-H, Juan C-H, Huang H-H, Chen L-F, Hsieh J-C et al. Efficacy of prefrontal theta-burst stimulation in refractory depression: a randomized sham-controlled study. Brain 2014; 137: 2088–2098.

6 Vanderwolf C. Hippocampal electrical activity and voluntary movement in the rat. Electroencephalogr Clin Neurophysiol 1969; 26: 407–418.

7 O’Keefe J, Recce ML. Phase relationship between hippocampal place units and the EEG theta rhythm. Hippocampus 1993; 3: 317–330.

8 Brazier MA. Studies of the EEG activity of limbic structures in man. Electroencephalogr Clin Neurophysiol 1968; 25: 309–318.

9 Chung SW, Sullivan CM, Rogasch NC, Hoy KE, Bailey NW, Cash RFH et al. The effects of individualised intermittent theta burst stimulation in the prefrontal cortex: A TMS-EEG study. Hum Brain Mapp 2018. doi:10.1002/hbm.24398.

10 Oberman L, Edwards D, Eldaief M, Pascual-Leone A. Safety of Theta Burst Transcranial Magnetic Stimulation: A Systematic Review of the Literature. J Clin Neurophysiol 2011; 28: 67–74.

11 Huang YZ, Edwards MJ, Rounis E, Bhatia KP, Rothwell JC. Theta burst stimulation of the human motor cortex. Neuron 2005; 45: 201–206.

12 Conte A, Rocchi L, Nardella A, Dispenza S, Scontrini A, Khan N et al. Theta-Burst Stimulation-Induced Plasticity over Primary Somatosensory Cortex Changes Somatosensory Temporal Discrimination in Healthy Humans. PLoS One 2012; 7: e32979.

13 Chung SW, Lewis BP, Rogasch NC, Saeki T, Thomson RH, Hoy KE et al. Demonstration of short-term plasticity in the dorsolateral prefrontal cortex with theta burst stimulation: A TMS-EEG study. Clin Neurophysiol 2017; 128: 1117–1126.

14 Anderson RJ, Hoy KE, Daskalakis ZJ, Fitzgerald PB. Repetitive transcranial magnetic stimulation for treatment resistant depression: Re-establishing connections. Clin Neurophysiol 2016; 127: 3394–3405.

15 Liston C, Chen AC, Zebley BD, Drysdale AT, Gordon R, Leuchter B et al. Default mode network mechanisms of transcranial magnetic stimulation in depression. Biol Psychiatry 2014; 76: 517–526.

16 Taylor SF, Ho SS, Abagis T, Angstadt M, Maixner DF, Welsh RC et al. Changes in brain connectivity during a sham-controlled, transcranial magnetic stimulation trial for depression. J Affect Disord 2018; 232: 143–151.

17 Mayberg HS. Targeted electrode-based modulation of neural circuits for depression. J Clin Invest 2009; 119: 717–725.

18 Weigand A, Horn A, Caballero R, Cooke D, Stern AP, Taylor SF et al. Prospective Validation That Subgenual Connectivity Predicts Antidepressant Efficacy of Transcranial Magnetic Stimulation Sites. Biol Psychiatry 2018; 84: 28–37.

19 Singh A, Erwin-Grabner T, Sutcliffe G, Antal A, Paulus W, Goya-Maldonado R. Personalized repetitive transcranial magnetic stimulation temporarily alters default mode network in healthy subjects. Sci Rep 2019; 9: 5631.

20 Greicius MD, Flores BH, Menon V, Glover GH, Solvason HB, Kenna H et al. Resting-State Functional Connectivity in Major Depression: Abnormally Increased Contributions from Subgenual Cingulate Cortex and Thalamus. Biol Psychiatry 2007; 62: 429–437.

21 Andreescu C, Tudorascu DL, Butters MA, Tamburo E, Patel M, Price J et al. Resting state functional connectivity and treatment response in late-life depression. Psychiatry Res Neuroimaging 2013; 214: 313–321.

22 Bluhm R, Williamson P, Lanius R, Théberge J, Densmore M, Bartha R et al. Resting state default-mode network connectivity in early depression using a seed region-of-interest analysis: Decreased connectivity with caudate nucleus. Psychiatry Clin Neurosci 2009; 63: 754–761.

23 Mulders PCP, van Eijndhoven PF, Schene AH, Beckmann CF, Tendolkar I. Resting-state functional connectivity in major depressive disorder: A review. Neurosci Biobehav Rev 2015; 56: 330–344.

24 Li B, Liu L, Friston KJ, Shen H, Wang L, Zeng L-L et al. A Treatment-Resistant Default Mode Subnetwork in Major Depression. Biol Psychiatry 2013; 74: 48–54.

25 Manoliu A, Meng C, Brandl F, Doll A, Tahmasian M, Scherr M et al. Insular dysfunction within the salience network is associated with severity of symptoms and aberrant inter-network connectivity in major depressive disorder. Front Hum Neurosci 2014; 7: 1–17.

26 Zhu X, Wang X, Xiao J, Liao J, Zhong M, Wang W et al. Evidence of a dissociation pattern in resting-state default mode network connectivity in first-episode, treatment-naive major depression patients. Biol Psychiatry 2012; 71: 611–617.

27 van Tol M-J, Li M, Metzger CD, Hailla N, Horn DI, Li W et al. Local cortical thinning links to resting-state disconnectivity in major depressive disorder. Psychol Med 2014; 44: 2053–2065.

28 Sheline YI, Price JL, Yan Z, Mintun MA. Resting-state functional MRI in depression unmasks increased connectivity between networks via the dorsal nexus. Proc Natl Acad Sci U S A 2010; 107: 11020–5.

29 Alexopoulos GS, Hoptman MJ, Kanellopoulos D, Murphy CF, Lim KO, Gunning FM. Functional connectivity in the cognitive control network and the default mode network in late-life depression. J Affect Disord 2012; 139: 56–65.

30 Berman MG, Peltier S, Nee DE, Kross E, Deldin PJ, Jonides J. Depression, rumination and the default network. Soc Cogn Affect Neurosci 2011; 6: 548–555.

31 Wu M, Andreescu C, Butters MA, Tamburo R, Reynolds CF, Aizenstein H. Default-mode network connectivity and white matter burden in late-life depression. Psychiatry Res Neuroimaging 2011; 194: 39–46.

32 Rossi S, Hallett M, Rossini PM, Pascual-Leone A, Avanzini G, Bestmann S et al. Safety, ethical considerations, and application guidelines for the use of transcranial magnetic stimulation in clinical practice and research. Clin Neurophysiol 2009; 120: 2008–2039.

33 Lefaucheur J-PP, Andre-Obadia N, Antal A, Ayache SS, Baeken C, Benninger DH et al. Evidence-based guidelines on the therapeutic use of repetitive transcranial magnetic stimulation (rTMS). Clin Neurophysiol 2014; 125: 2150–2206.

34 Oldfield RC. The assessment and analysis of handedness: The Edinburgh inventory. Neuropsychologia 1971; 9: 97–113.

35 Liu Q, Zhou R, Chen S, Tan C. Effects of Head-Down Bed Rest on the Executive Functions and Emotional Response. PLoS One 2012; 7. doi:10.1371/journal.pone.0052160.

36 Moeller S, Yacoub E, Olman CA, Auerbach E, Strupp J, Harel N et al. Multiband multislice GE-EPI at 7 tesla, with 16-fold acceleration using partial parallel imaging with application to high spatial and temporal whole-brain fMRI. Magn Reson Med 2010; 63: 1144–1153.

37 Setsompop K, Gagoski BA, Polimeni JR, Witzel T, Wedeen VJ, Wald LL. Blipped-controlled aliasing in parallel imaging for simultaneous multislice echo planar imaging with reduced g-factor penalty. Magn Reson Med 2012; 67: 1210–1224.

38 Jenkinson M, Beckmann CF, Behrens TEJ, Woolrich MW, Smith SM. FSL. Neuroimage 2012; 62: 782–790.

39 Brett M, Anton J-L, Valbregue R, Poline J-B. Region of interest analysis using an SPM toolbox. In: NeuroImage Academic Press, 2002, pp 769–1198.

40 Kaiser RH, Andrews-Hanna JR, Wager TD, Pizzagalli DA. Large-Scale Network Dysfunction in Major Depressive Disorder. JAMA Psychiatry 2015; 02478: 603–611.

41 Fox MD, Buckner RL, White MP, Greicius MD, Pascual-Leone A. Efficacy of transcranial magnetic stimulation targets for depression is related to intrinsic functional connectivity with the subgenual cingulate. Biol Psychiatry 2012; 72: 595–603.

42 Menon V. Large-scale brain networks and psychopathology: a unifying triple network model. Trends Cogn Sci 2011; 15: 483–506.

43 Menon V, Uddin LQ. Saliency, switching, attention and control: a network model of insula function. Brain Struct Funct 2010; 214: 655–667.

44 Fox MD, Raichle ME. Spontaneous fluctuations in brain activity observed with functional magnetic resonance imaging. Nat Rev Neurosci 2007; 8: 700–711.

45 Hamilton JP, Furman DJ, Chang C, Thomason ME, Dennis E, Gotlib IH. Default-Mode and Task-Positive Network Activity in Major Depressive Disorder: Implications for Adaptive and Maladaptive Rumination. Biol Psychiatry 2011; 70: 327–333.

46 Sridharan D, Levitin DJ, Menon V. A critical role for the right fronto-insular cortex in switching between central-executive and default-mode networks. Proc Natl Acad Sci U S A 2008; 105: 12569–12574.

47 Iwabuchi SJ, Peng D, Fang Y, Jiang K, Liddle EB, Liddle PF et al. Alterations in effective connectivity anchored on the insula in major depressive disorder. Eur Neuropsychopharmacol 2014; 24: 1784–1792.

48 Cooney RE, Joormann J, Eugène F, Dennis EL, Gotlib IH. Neural correlates of rumination in depression. Cogn Affect Behav Neurosci 2010; 10: 470–478.

49 Iwabuchi SJ, Auer DP, Lankappa ST, Palaniyappan L. Baseline effective connectivity predicts response to repetitive transcranial magnetic stimulation in patients with treatment-resistant depression. Eur Neuropsychopharmacol 2019;: 1–10.

50 Baeken C, Duprat R, Wu G-R, De Raedt R, van Heeringen K. Subgenual Anterior Cingulate– Medial Orbitofrontal Functional Connectivity in Medication-Resistant Major Depression: A Neurobiological Marker for Accelerated Intermittent Theta Burst Stimulation Treatment? Biol Psychiatry Cogn Neurosci Neuroimaging 2017; 2: 556–565.

51 Duprat R, Wu GR, De Raedt R, Baeken C. Accelerated iTBS treatment in depressed patients differentially modulates reward system activity based on anhedonia. World J Biol Psychiatry 2017; 0: 1–12.

