## Supplementary Information for "Intermittent theta burst stimulation at personalized targets reduces the functional connectivity of the default mode network in healthy subjects"

### Affiliations:

**Supplementary Materials**

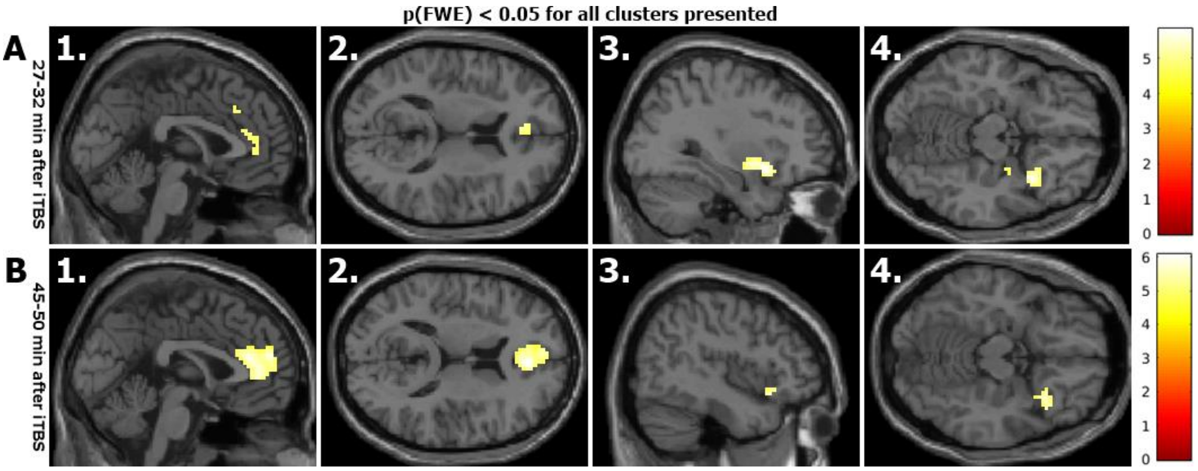


Supplementary Figure 1: Changes in functional connectivity of default mode network (DMN) after real iTBS without comparison to sham condition. The effects of real iTBS (without sham comparison) are very similar to those obtained when real iTBS is compared against sham iTBS (Figure 3), except by smaller mPFC and larger rAI blobs in the R2 rsfMRI (Suppl. Figure 1 – A1-A4).

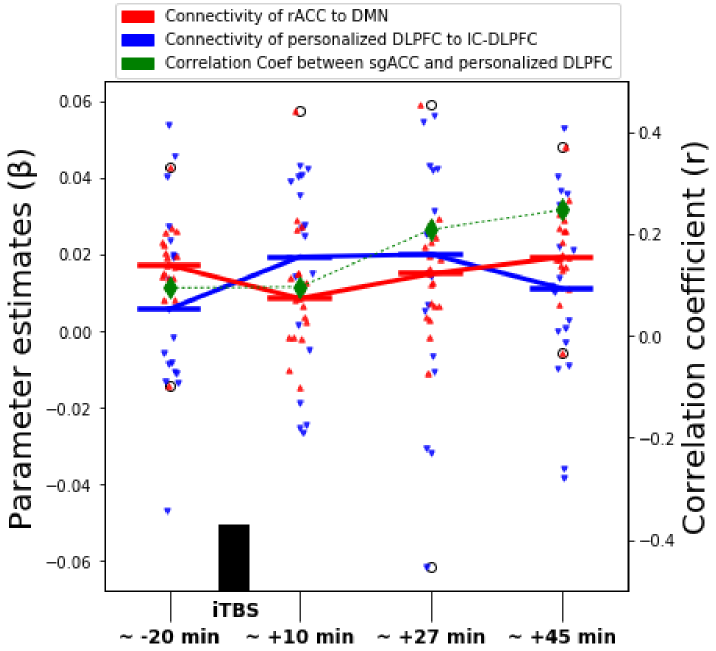


Supplementary Figure 2: Left axis shows the parameter estimates of left DLPFC (blue) and rACC (red) of IC-DLPFC and DMN, respectively. Dots represent the individuals and horizontal lines depict the median of the parameter estimates for the respective rsfMRI window. Only minor changes in the median of parameter estimates ranging between (0-0.02) and correlation coefficient between (0-0.17) are observed. This implies the functional connectivity fluctuates around the baseline during all rsfMRI sessions and the interaction between sgACC and personalized left DLPFC is also not affected by sham iTBS.
